# Does crossed aphasia originate from developmental disorders? A Mini-review and case study

**DOI:** 10.1101/039024

**Authors:** Anna B. Jones, Thomas H Bak, Mark E. Bastin, Joanna M. Wardlaw, Cyril R. Pernet

## Abstract

Cognitive impairments associated with crossed aphasia were investigated in a single case study and a review of the literature. A review of literature identifies 4 main cognitive co-morbidities that are significantly associated with crossed aphasia. We present a case of confirmed crossed aphasia with dyslexia and dysgraphia, in which the latter two cannot be fully explained by the current lesion and are probable developmental disorders (dyslexia/dysgraphia). Extensive longitudinal cognitive investigations and a series of advanced imaging techniques (structural and functional) were used to investigate the cognitive and neuroanatomical basis of crossed aphasia and associated impairments in this patient. Using the results from the literature review and the single case study, we suggest that developmental disorders can be an underlying cause of partial right lateralisation shift of language processes, thereby supporting the theory that developmental disorders can be an underlying cause of crossed aphasia.

**Highlights:**

- Central apraxia, dysgraphia, hemi-neglect & acalculia associated with CA
- Developmental disorders can underlie partial right lateralisation shift
- Dysfunction of left hemisphere can cause crossed aphasia
- Clinically, pre-morbid impairments must be investigated in CA cases

## 1. Introduction

In 1863, Paul Broca (1863) described a strict anatomo-functional connection between the handedness of a patient (i.e. right dominant motor control) and the hemispheric control over language functions. Twenty-eight years later, Oppenheim (1891) presented two cases of right-handed individuals who suffered right hemisphere lesions and subsequent aphasia, thus questioning the hypothesis of motor-language co-dominance. Following these observations, Bramwell (1899) proposed the term ‘Crossed Aphasia’ to denote aphasia caused by an ipsilateral lesion to the dominant hand. Wada and Rasmussen (1960) later showed that ∽70% of left-handed patients also have left hemispheric dominance for language, thereby supporting Goodglass and Quadfasel's study (1954) suggesting that crossed aphasia occurs in 70-80% of left-handed cases and therefore the term ‘Crossed Aphasia’ (CA) became a synonymous term for ‘Crossed Aphasia in Dextrals’ (CAD). Specific criteria for a diagnosis of ‘crossed aphasia’ were first put in place by Brown and Wilson (1973) and reviewed by Habib et al. (1983) and Coppens and Robbey (1992). Mariën et al. (2004) conducted a review of the CA literature and concluded an algorithm of diagnostic criteria for vascular CA in adults widely in use today. This algorithm divides patients into unreliable, possible and reliable CA cases. Patients are only classed as reliable CA cases if they have 1. clear-cut evidence of language disorder, right-handedness and morphological integrity of the left hemisphere; as well as 2. absence of left-handedness in relatives and/or early brain damage/seizures in infancy. To establish a clear-cut language disorder, and further understand the underlying neural substrates associated with CA, it is important to investigate co-morbidities (Marien, Engelborghs, Vignolo, & De Deyn, 2001). Co-morbidities are already widely acknowledged in the literature but are seldom investigated beyond the level of diagnosis. Dysgraphia for instance is frequently reported in dextral patients with aphasia from a left-hemisphere lesion (for example: Roeltgen & Heilman, 1984; Sinanovic, Mrkonjic, Zukic, Vidovic, & Imamovic, 2011; Tanridag & Kirshner, 1985), but there are only a few reports of patients with confirmed CA exhibiting dysgraphia (Assal, Perentes, & Deruaz, 1981; Marien, Engelborghs, Vignolo, & De Deyn, 2001; Mastronardi et al., 1994) and no known reports of anomic aphasia with persistent dysgraphia. Specific lesion localisation, anatomical mapping and network disruption within CA has not been studied extensively either. Kim et al.'s review (2013) found that the regions most frequently involved in CA are the right lentiform nucleus, in particular the putamen, and basal ganglia. The right parahippocampal gyrus, claustrum, frontal lobe and precentral gyrus were also locations found to be involved in CA (Kim, Yang, & Paik, 2013).

There are four main hypotheses for the possible causes underlying CA (Cappa et al., 1993): 1. dysfunction within the left hemisphere, either congenital or acquired, causing a lateralisation shift (Bakar, Kirshner, & Wertz, 1996; Bhatnagar, Imes, Buckingham, & Puglishi-Creegan, 2006; Cappa et al., 1993). 2. Bilateral representation of language functions 3. Genetic basis (Alexander & Annett, 1996; Osmon, Panos, Kautz, & Gandhavadi, 1998) - i.e. the ‘right-shift’ (RS) theory (Annett, 1985). Cohen et al. (1993) also suggest a genetic underpinning, but in relation to the co-existence of anomalous cerebral language dominance and situs inversus. 4. Diaschisis, i.e. language areas in the left hemisphere are functionally depressed indirectly as a result of functional connections between the lesion in the right hemisphere and the preserved areas (Finger, Koehler, & Jagella, 2004). Here, we performed a review of co-morbidities associated with CA to test these hypotheses. In particular, if other deficits of known lateralized cognitive processes are associated to CA, this will allow the rejection of hypotheses 1 (only auditory and/or spoken language if right lateralized) and 2 (all language functions are bilateral) and further distinguish between hypotheses 3 and 4 (true right lateralization versus diachisis). We then detail a new case of CA in a right-handed English speaking male (CF) who suffered from an acute right middle cerebral artery infarct, and discuss his deficits in light of the results obtained in the review.

## 2. Materials and Methods

### 2.1 Mini-review

A search of the literature was conducting using the following parameters in PubMed and ScienceDirect: ((crossed aphasi*[Title]) OR (crossed dysphasi*[Title]) OR ((right hemisphere stroke[Title]) AND (aphasi*[Title])) OR ((right hemisphere stroke[Title]) AND (dysphasi*[Title]))). 82 articles were retrieved, and using Marien et al.'s algorithm (Marien, Paghera, De Deyn, & Vignolo, 2004) to select only reliable cases of CA, 23 articles were selected. A further 7 articles were retrieved from cited references, totalling 30 papers with 57 ‘reliable’ CA cases. After listing all co-morbidities, the 6 most common were selected and the number of cases was adjusted for the number of cases tested (e.g. a disorder might be present in all patients, but we don’t know about it because not tested). The adjustment was computed as the number of cases presenting a given deficits * total number of CA cases/number of patients tested for that deficit. As there is no current epidemiological data available for the cognitive co-morbidities identified with crossed aphasia, a simple binomial model that assumes equal probability of occurrence for each condition was used (i.e. occurrence above 33%, corrected for multiple comparisons, upper bound of the 95% confidence intervals, is considered significant).

Among all co-morbidities, apraxia was split into “Central” (i.e. apraxia caused by impaired initiation e.g. constructional and ideomotor) and “Peripheral” (i.e. apraxia caused by impaired execution e.g. limb and oral) and only “central” apraxia was included since “peripheral” apraxia is not related to the language system. Visuo-spatial deficits were classified separately to visual-field neglect (a neuropsychological condition causing a deficit in attention to, and awareness of, one side of space) as they often occurred separately and/or were not both assessed. Visuo-spatial impairments observed across studies corresponded mainly to difficulties with visual organisation, spatial relations and position discrimination, constructive abilities, visual memory and visual scanning speed. Due to the heterogeneous nature of these impairments and the assessment procedures used in the studies included, visuo-spatial impairments were not included as one of the main cognitive co-morbidities in this review.

### 2.2 Case study

*Ethics Statement:* This study was approved by the NHS Lothian South East Scotland Research Ethics Committee 01 (Research Ethics Committee reference number: 11/SS/0055). Full information was provided to the participant, in an aphasia-friendly format and full informed consent was obtained from the participants.

#### 2.2.1 Diagnostic

To confirm the presence of crossed aphasia the integrity of the left hemisphere was investigated on CT (acute phase) and MRI (chronic phase - T1 3D IRP). Handedness was assessed using interview and the Edinburgh Handedness Inventory test (Oldfield, 1971). Aphasia was assessed using the Western Aphasia Battery (WAB) (Kertesz, 1982) at three separate time points: acute (1-6 weeks post stroke), sub-acute (2-6 months post stroke) and chronic (7-11 months post stroke).

#### 2.2.2 Cognitive Assessment

Neuropsychological testing was conducted to assess CF's language, memory, visuo-spatial skills, executive functioning, calculia (Ardila & Rosselli, 2002; Leff et al., 2009) and retention of specific knowledge relate to his pre-morbid field of expertise (Graham, Lambon Ralph, & Hodges, 1999; Graham, Patterson, Pratt, & Hodges, 1999; Omar, Hailstone, Warren, Crutch, & Warren, 2010; Robinson, Rossor, & Cipolotti, 1999).

To explore the exact breakdown of CF's language, assessments were carried out using subtests of the Psycholinguistic Assessment of Language Processing in Aphasia (PALPA, Kay, Lesser, & Coltheart, 1992); and an experimental battery of semantic assessments and an informal auditory phonological processing assessment (Table A.6). Semantic assessments consisted of The Pyramids and Palmtrees Test (PPT, Howard, 1992); Kissing and Dancing Test (KDT, Bak & Hodges, 2003); Tomato and Tuna Test (TTT, Faber et al., 2008); and Sound to Picture Matching Test (SPMT, Bozeat, Lambon Ralph, Patterson, Garrard, & Hodges, 2000). All of these assessments were shortened, adapted and complied for experimental purposes and programmed to run in E-prime 2.0 (Schneider, Eschman, & Zuccolotto, 2012) on a laptop computer. The raw score gained from each of these assessments was the number of errors made.

Memory was assessed testing verbal and non-verbal immediate recall, delayed recall, and processing speed [Doors subtest of the Doors and People Assessment, Brit Memory and Information Processing Battery, Rey-Osterrieth Complex Figure Test, Digit Span subtest of the Weschler Adult Intelligence Scale-IV, Weschler Memory Scales III] (Baddeley, Emslie, & Nimmo-Smith, 1994; Coughlan, Oddy, & Crawford, 2007; Fastenau, Denburg, & Hufford, 1999; Wechsler, 2008; Wechsler, 1999). Visuo-spatial skills were assessed using subsections of the Visual Object & Space Perception Battery (Lezak, Howieson D., Loring D., Hannay H., & Fischer J., 2004) as well as the ‘direct copy’ subsection from the Rey-Osterrieth Complex Figure task (Fastenau, Denburg, & Hufford, 1999). Executive Functioning was investigated using the matrix reasoning subtest from the WAIS IV (Wechsler, 2008) and the Key Search, Temporal Judgement, and Modified 6 elements subtests from the Behavioural Assessment of Dysexecutive Function battery (Wilson, Alderman, Burgess, Emslie, & Evans, 1996). Acalculia was informally tested using the following mathematical tasks: addition, subtraction and deciding which is the larger/smaller out of two numbers (Table A.1). Premorbid semantic knowledge was investigated using an informal questionnaire (Figure A.1).

Standardised assessment results were compared against published normative data and performance descriptors (impaired/unimpaired) applied. If published normative data was unavailable results were compared against published control data. An impaired performance descriptor was applied to all scores below 100% on informal assessments without published control data. Results were mapped onto an adapted and extended cognitive-neuropsychological model of single word processing (Ellis et al., (1988), Ellis (1998) Laganaro and Alario, 2006; Martin et al., 1999; Nickels et al., 1997 -*Figure 4*) with the aim of identifying the nature of the underlying impairments. Assessments were matched to the model according to the different processing components used within the task. Performance descriptors, error analysis and convergent evidence from different assessments were then used to identify if a processing component within the module was intact or impaired (Whitworth et al. 2005).

#### 2.2.3 Imaging

In addition to the CT scan performed at the acute time point, high-resolution structural, DTI and fMRI data were obtained at the chronic phase (75 weeks post-stroke onset) using a GE Signa HDxt 1.5 T clinical scanner at the Brain Research Imaging Centre (http://www.bric.ed.ac.uk), University of Edinburgh, UK. <u>Structural imaging consisted of the following sequences:</u> (i) a T1-weighted volume (3D IRP - 180 slices, 2 mm thick coronal slices, 1.3 x 1.3 mm in-plane resolution with a 256 mm FOV); (ii) a T2-weighted volume (FSE - 72 slices, 2 mm thick axial slices thickness, 1 x 1 mm in-plane resolution with a 256 mm FOV), and (iii) a FLAIR-weighted volume (FSE - 40 slices, 4mm thick axial slices, 1 x 1.3 mm in-plane resolution with a 256 mm FOV). The DTI examination consisted of 7 T2-weighted (b = 0 s mm-2) and sets of diffusion-weighted (b = 1000 s mm-2) single-shot spin-echo echo-planar imaging (EPI) volumes acquired with diffusion gradients applied in 64 non-collinear directions. Volumes were acquired in the axial plane, with a FOV of 256 x 256 mm, 72 contiguous slice locations, and image matrix and slice thickness designed to give 2 mm isotropic voxels. The repetition and echo time for each EP volume were 16.5 s and 98 ms respectively. fMRI was also performed to map language areas. The auditory cortex and Wernicke area were mapped using a word repetition task whilst Broca's area was mapped using a verb generation task (Gorgolewski, Storkey, Bastin, Whittle, & Pernet, 2013), and the Visual Word Form Area was mapped using a one-back visual detection task with 8 blocks of 16 sec per category, showing checkerboards, faces (from ‘Labelled Faces in the Wild’, http://vis-www.cs.umass.edu/lfw/index.html), objects (from the Amsterdam Library of Object Images, (Geusebroek, Burghouts, & Smeulders, 2005) and the Department of Cognitive, Linguistic & Psychological Sciences, Brown University (http://titan.cog.brown.edu:8080/TarrLab) and high frequency nouns from http://www.esldesk.com/esl-quizzes/frequently-used-english-words/words.htm) (Cohen et al., 2000b; Dehaene & Cohen, 2011; Price & Devlin, 2011). fMRI data was acquiring as follows: a) *Word repetition task* and b) *Verb generation task* - (EPI - 30 slices, 4mm thick axial slices, 4 x 4 mm in-plane resolution with a 256 mm FOV); c) *Passive visual word form task -* (EPI - 27 slices, 4 mm thick axial slices, 4 x 4 mm in-plane resolution with a 256 mm FOV).

#### 2.2.4 Tract-based Spatial Statistics

All DTI data were converted from DICOM (http://dicom.nema.org) to NIfTI-1 (http://nifti.nimh.nih.gov/nifti-1) format using the TractoR package for fibre tracking analysis (http://www.tractor-mri.org.uk). FSL tools (http://www.fmrib.ox.ac.uk/fsl) were then used to extract the brain, remove bulk motion and eddy current induced distortions by registering all subsequent volumes to the first T2-weighted EPI volume, estimate the water diffusion tensor and calculate parametric maps of MD and FA from its eigenvalues using DTIFIT.

Following protocols described in detail by ENIGMA (Enhancing Neuro Imaging Genetics Through Meta Analysis; http://enigma.ini.usc.edu/protocols/dti-protocols/#eDTI), differences in CF's MD values in language tracts (left and right internal capsule, inferior and superior longitudinal fasciculi and splenium of corpus callosum) were compared with 5 aged matched (63.3 ± 1.4 years) healthy controls assessed in regions-of-interest (ROI) extracted from white matter skeletons produced using Tract-based Spatial Statistics (TBSS; http://www.fmrib.ox.ac.uk/fsl). First, all FA volumes were linearly and non-linearly registered to the standard FMRIB58_FA volume. Second, a white matter skeleton was created from the mean of all registered FA volumes. This was achieved by searching for maximum FA values in directions perpendicular to the local tract direction in the mean FA volume. An FA threshold of 0.25 was applied to the skeleton to exclude predominantly non-white matter voxels. Third, for each subject's FA volume, the maximum voxel perpendicular to the local skeleton direction was projected onto the skeleton. This resulted in one FA skeleton volume per subject corresponding to centres of white matter structures. Average MD values were then obtained for the six subjects from skeletal projections in language tract ROI defined using FSL's JHU white matter atlas. For each ROI 95% percentile bootstrapped confidence intervals of the trimmed mean, (adjusted for multiple testing) were computed and compared to CF's values (see Figure A.2). A lateralisation index (left-right/left+right) was applied to MD values for the specified language tracts for the 5 aged matched healthy controls (and percentile bootstrapped CI calculated) and compared to CF (Lebel and Beaulieu, 2009) (see Figure A.2 and Table A.2).

#### 2.2.5 fMRI data analysis

For each fMRI task SPM12 was used. Data were first slice-time corrected (amount of correction varied according to specific task parameters, but in all cases the data was temporally aligned to the middle temporal slice), then realigned to the 1^st^ image of each session and then to the mean EPI (SPM12 default parameters) and finally smoothed at 6mm isotropic Gaussian kernel. The T1 image (following the structural processing above) was then co-registered onto the mean EPI and the transformation parameters applied to the gray matter mask created earlier. The General Linear Model (Friston et al., 1994) was used to estimate the BOLD signal response for each task separately with parameter estimates restricted to the gray matter mask. For each task, one regressor per condition was used (1 regressor for activation blocks in the word repetition; 1 regressor for activation blocks in the verb generation task; 4 regressors for the blocks of faces/ objects/ words/ checkerboards in the passive visual work form task) as well as motion parameters and motion outlier censoring (Siegel et al., 2014). Adaptive thresholding (Gorgolewski, Storkey, Bastin, & Pernet, 2012) was used to obtain the single subject statistical maps for each task.

Hemispheric lateralisation of the whole brain and the temporal lobe for the Word repetition task (a) and the Verb generation task (b) was assessed, based on number of activated voxels, using the LI-tool (Wilke & Lidzba, 2007) in SPM12, producing lateralisation indices (LI) in each task (LI>0 signifies left hemispheric lateralisation, LI>0 signifies right hemispheric lateralisation). To allow for comparison, fMRI data from 10 healthy controls participants (median age 52.5 years, 3 left-handed and 7 right-handed) (Gorgolewski et al., 2013) who had carried out the same two tasks was also analysed for hemispheric lateralisation of the whole brain and temporal lobe, using the LI-tool (Wilke & Lidzba, 2007).

## 3. Results

### 3.1 Mini-review

Using Marien et al.'s algorithm (Marien, Paghera, De Deyn, & Vignolo, 2004) 57 ‘reliable’ CA cases were identified, 52 as a result of stroke and 5 as a result of a tumour (Alexander & Annett, 1996; Habib, Joanette, Ali Cherif, & Poncet, 1983; April & Han, 1980; Assal, Buttet, & Jolivet, 1981; Bakar, Kirshner, & Wertz, 1996; Bartha, Marien, Poewe, & Benke, 2004; Bhatnagar, Imes, Buckingham, & Puglishi-Creegan, 2006; Bhatnagar, Buckingham, Puglisi-Creegan, & Hacein-Bey, 2011; Cappa et al., 1993; Cohen, Grony, Hermine, Gray, & Degos, 1993; De Witte, Verhoeven, Engelborghs, De Deyn, & Marien, 2008; Denes & Caviezel, 1981; Faglia & Vignolo, 1990; Giovagnoli, 1993; Ha, Pyun, Hwang, & Sim, 2012; Haaland & Miranda, 1982; Habib, Joanette, Ali-Cherif, & Poncet, 1983; Henderson, 1983; Ishizaki et al., 2012; Kim, Yang, & Paik, 2013; Lessa Mansur, Radanovic, Santos Penha, Iracema Zanotto de Mendonoa, & Cristina Adda, 2006; Marien, Engelborghs, Vignolo, & De Deyn, 2001; Marshall & Halligan, 1992; Mastronardi et al., 1994; Osmon, Panos, Kautz, & Gandhavadi, 1998; Paghera, Marin, & Vignolo, 2003; Paparounas, Eftaxias, & Akritidis, 2002; Patidar et al., 2013; Rey, Levin, Rodas, Bowen, & Nedd, 1994; Stefanis, Desmond, & Tatemichi, 1997). Across those studies, six main co-morbidities were found: Central Apraxia, Dysgraphia, Hemi-neglect, Acalculia, Attentional deficits, Memory deficits (see Figure 1 for a full breakdown). Under the hypothesis they all have equal chance to co-occur (chance level: 6 to 33%), central apraxia, dysgraphia, left visual field neglect and acalculia were found to significantly co-occur with CA (*Table 1*). (See Table A.3 for a detailed review of studies involved).

**Table 1:** Breakdown of co-morbidities detailed in the literature, showing the total number of cases assessed and the associated number of cases identified. The adjusted number of cases is the estimated number of cases out of the 57 unique cases of confirmed CA. Italics signify cognitive comorbidities that significantly co-occur with crossed aphasia.

| **Cognitive co-morbidity** | **Number of cases tested** | **Number of cases identified** | **Adjusted number of cases** |
| --- | --- | --- | --- |
| *Central Apraxia* | 31 | 21 | 39 |
| *Dysgraphia* | 36 | 22 | 35 |
| *Left VF neglect* | 33 | 22 | 38 |
| *Acalculia* | 18 | 13 | 42 |
| Attention | 14 | 4 | 15 |
| Memory | 15 | 5 | 19 |

**Figure 1:**
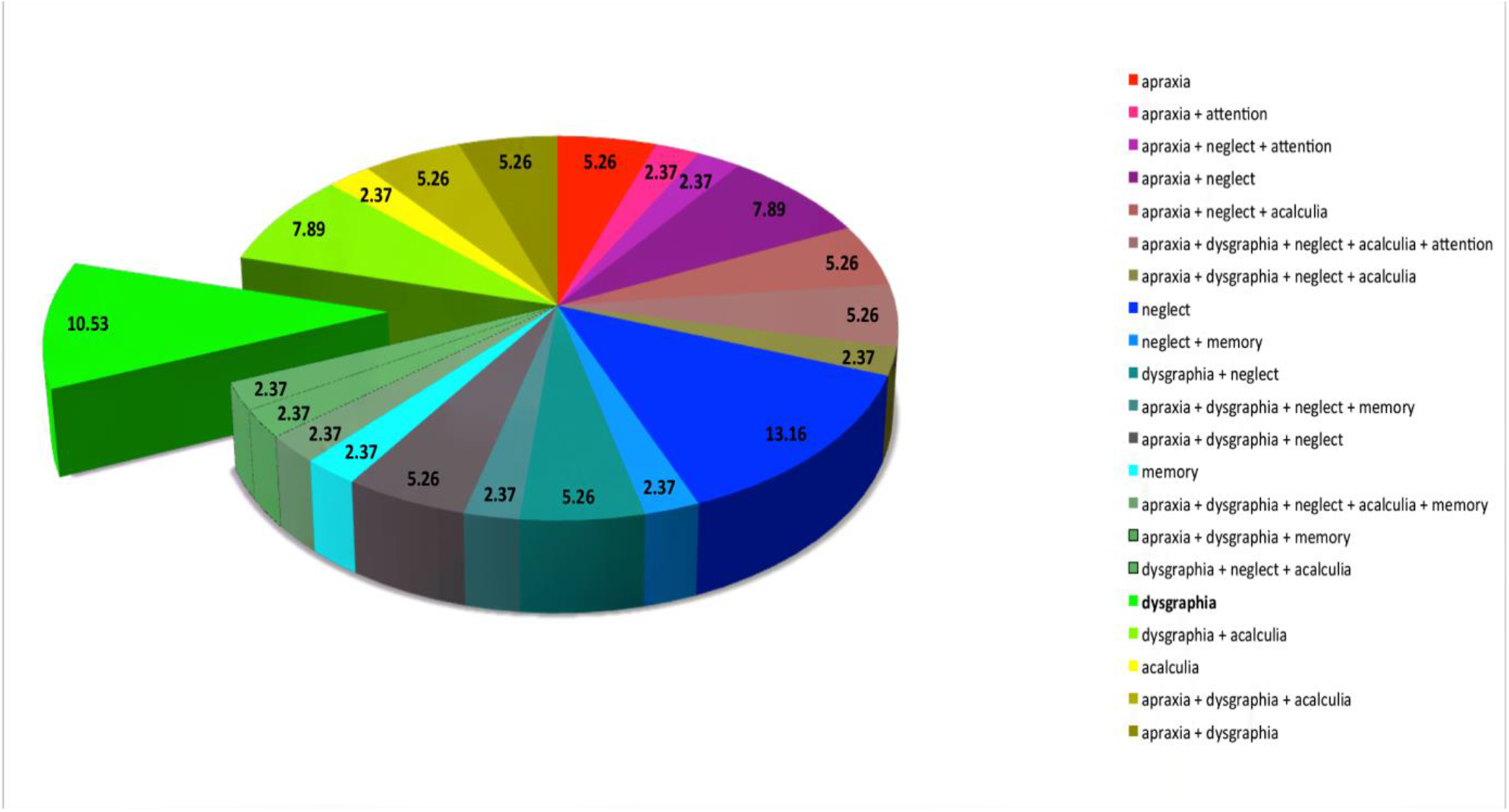
Pie chart showing occurrence of the 6 main coggnitive co-morbidities present alongside confirmed cases of Crossed Aphasia in the literature. Percentages correspond to the number of times one or more cognitive impairment co-occurred (total number of classes identified =21) / total number of confirmed CA cases with at least one of the main co-morbidities present (n=38). The exploded pie segment (dysgraphia) is the class of co-morbidities corresponding with CF's symptomatology at the chronic stage.

Aphasia and apraxia are two independent conditions, but they are often associated. The language and praxis systems share a number of functional features (i.e. sensory-motor integration and symbolic representation) and rely on common anatomical structures involving the frontal cortex and basal ganglia (Gross & Grossman, 2008; Kobayashi & Ugawa, 2013). Double dissociations have however been found between aphasia and apraxia (Papagno, la Sala, & Basso, 1993), suggesting that the two networks do not completely overlap. Praxis relies on a large-scale network involving frontal-parietal-regions and basal ganglia. Central apraxia is commonly associated with right hemispheric damage, and is most likely caused by a higher-order visuospatial processing deficit in patients with parietal (intraparietal sulcus) lesions, or by impairments to organisation and planning in patients with frontal (middle frontal gyrus - MFG) lobe damage (Haaland, Harrington, & Knight, 2000). As both the intraparietal sulcus and the MFG are involved in language (syntax (Carreiras, Carr, Barber, & Hernandez, 2010), dorsal language pathway (Herman, Houde, Vinogradov, & Nagarajan, 2013) and phoneme detection (Simon, Mangin, Cohen, Le Bihan, & Dehaene, 2002), lexical-syntactic retrieval (Acheson & Hagoort, 2013), respectively), it stands to reason that central apraxia significantly co-occurs with CA, if we consider that only language is inversely lateralized in those individuals.

Dysgraphia impairments observed across studies corresponded to misspelling of words; neologisms, paraphasias and jargon; perseveration of letters or words; semantic errors; syntactical errors; spatial problems; morphological errors; word or letter omissions, substitutions, additions and deletions; and motor difficulties. Central writing processes are defined as ‘the retrieval of abstract orthographic word-forms, via orthographic lexicon or phoneme-to-grapheme conversion mechanisms, and their temporary storage in the graphemic buffer’ (Planton, Jucla, Roux, & Demonet, 2013) (p. 2773). Note that motor and linguistic impairments involved in writing are often associated with other abilities (i.e. praxis, literacy etc.) and thus not included in this definition. For the purposes of this study only those impairments involving ‘writing specific’ processes (i.e. orthographic coding - not including syntax, semantics, spatial or motor difficulties) were classified as ‘dysgraphic’. Two out of the three major anatomical regions involved in the writing system are also damaged in some aphasias: the superior frontal gyrus (SFG) involved in the recollection of grapheme representations and the supramarginal gyrus (SMG) involved in phoneme-to-grapheme conversion (Planton, Jucla, Roux, & Demonet, 2013). Under the constrictive criteria discussed above, dysgraphia significantly co-occurs with crossed aphasia, which is expected if one assumes that all language areas (i.e. not just perisylvian ones) are right lateralized.

Unilateral visual-field neglect is a common neurological presentation predominantly following damage to the right ventral fronto-parietal cortex i.e. a proposed distributed ventral attention network involving right frontal, temporal and parietal cortex (Corbetta & Shulman, 2011), which in turn disrupts the dorsal attention network. Importantly the dorsal attention network can be impaired (causing neglect symptoms) by damage to a variety of right hemispheric ventral fronto-parietal regions (Corbetta & Shulman, 2011). The data presented here showing that left visual-field neglect is significantly associated with CA are supportive of both the right hemisphere dominance of the visual attention system and also the wide variety of fronto-parietal regions corresponding with neglect. It has been postulated that these regions vary in terms of criticality to the overall attention process and network, with the more posterior regions more crucially involved (Callejas, Shulman, & Corbetta, 2014), specifically the inferior parietal lobule, temporo-parietal junction, the superior temporal lobe and the angular gyrus (Mort et al., 2003; Karnath, Berger, Kuker, & Rorden, 2004; Karnath & Rorden, 2012; Gillebert et al., 2011). In cases of crossed aphasia left visual field neglect commonly occurs alongside central apraxia, which again is expected if language only is inversely lateralized in those individuals.

Acalculia significantly co-occurs alongside crossed aphasia with the highest prevalence overall. Numerical cognition consists of a fronto-parietal network involving intra-parietal and pre-frontal areas (Moeller, Willmes, & Klein, 2015). The triple-code model of numerical processing (Dehaene, Piazza, Pinel, & Cohen, 2003) details three circuits (visual system encoding Arabic numbers; quantity system encoding analogical-semantic representations of size and distance relations; verbal system encoding numerals lexically, phonologically and syntactically) co-existing in the parietal lobe, specifically in the bilateral superior parietal gyrus, bilateral intra-parietal gyrus and the left angular gyrus. The frontal section of the pathway involves the pre-frontal cortex, in particular the inferior, medial and superior frontal gyri (Simon, Mangin, Cohen, Le Bihan, & Dehaene, 2002). Fronto-parietal association fibres (superior longitudinal fasciculus dorsally and external capsule ventrally) are also involved in numerical cognition (Moeller, Willmes, & Klein, 2015). Therefore, lesions in a number of parietal regions can be attributed to both language and numeracy impairments, for example the intraparietal sulcus is involved in number processing/arithmetic calculations (Seghier, Ramlackhansingh, Crinion, Leff, & Price, 2008) as well as numerous components of language processing [syntax processing (Carreiras, Carr, Barber, & Hernandez, 2010), phoneme detection (Simon, Mangin, Cohen, Le Bihan, & Dehaene, 2002) and the dorsal language pathway (Herman, Houde, Vinogradov, & Nagarajan, 2013)]. The significant cooccurrence of acalculia is expected as a consequence of the disruption between the quantity and the verbal systems, only this one being right rather than left lateralized.

### 3.2 Case study

CF was 66 year-old monolingual, right-handed English native speaker male laterality quotient (LQ) = +100, Decline R.10 (Oldfield, 1971)) admitted to hospital with a left sided weakness (including a left sided facial droop), confusion, apraxia and aphasia. A clinical CT scan showed an acute right middle cerebral artery (MCA) infarct. After 5 days in an acute ward, he was transferred to a stroke rehabilitation unit where he received speech and language therapy (SLT) for 1 month. He was then discharged home and immediately received SLT weekly for the following 11 months. All acute motor impairments resolved prior to transfer to inpatient rehabilitation; however, CF was left with a mixed communication impairment (aphasia and dysgraphia). CF was high functioning in all aspects of his life prior to his stroke. CF noted that he had difficulties as a child with written spelling and had suspected developmental dysgraphia. No other communication or visual problems were present before CF's admission for this episode. CF had a bilateral, symmetrical sensorineural hearing loss, corrected by bilateral hearing aids. At the time of testing, he showed mild low mood but did not have clinical depression (Depression Intensity Scale Circles, (DISCS, Turner-Stokes, Kalmus, Hirani, & Clegg, 2005) and Visual Analogue Self-Esteem Scale, (VASES, Brumfitt & Sheeran, 1999)).

#### 3.2.1 Lesion location

A CT scan (acute phase), confirmed by an MRI scan (chronic phase), showed complete integrity of the left hemisphere and revealed in the right hemisphere: (i) low attenuation and loss of grey/white matter differentiation, with mild swelling, in the insula, internal capsule, frontal operculum, and part of the inferior frontal gyrus and the mid frontal gyrus, (ii) a loss of basal ganglia definition, (iii) a mild degree of mass effect associated with the ischaemic lesion, and (iv) hyperdense MCA at the level of bifurcation and proximal M2 branches, signifying an intravascular thrombus as the cause of the infarct. Examination of fractional anisotropy (FA) maps obtained from diffusion tensor MRI (DTI) also suggests alterations in fibre structure around the internal capsule and arcuate branch of the superior frontal fasciculus. A statistical comparison of mean diffusivity (MD) values of CFs’ language tracts against 5 aged matched healthy controls not only confirmed reductions in structural integrity of the anterior, posterior and retro-splenial limb of the internal capsule and of the superior frontal fasciculus (*Figure 2*) but also defects in the right inferior frontal fasciculus and bilateral uncinate fasiculi (see *Table A.2)*. For these last two tracks, both left and right MD values were much larger, than in the controls.

**Figure 2:**
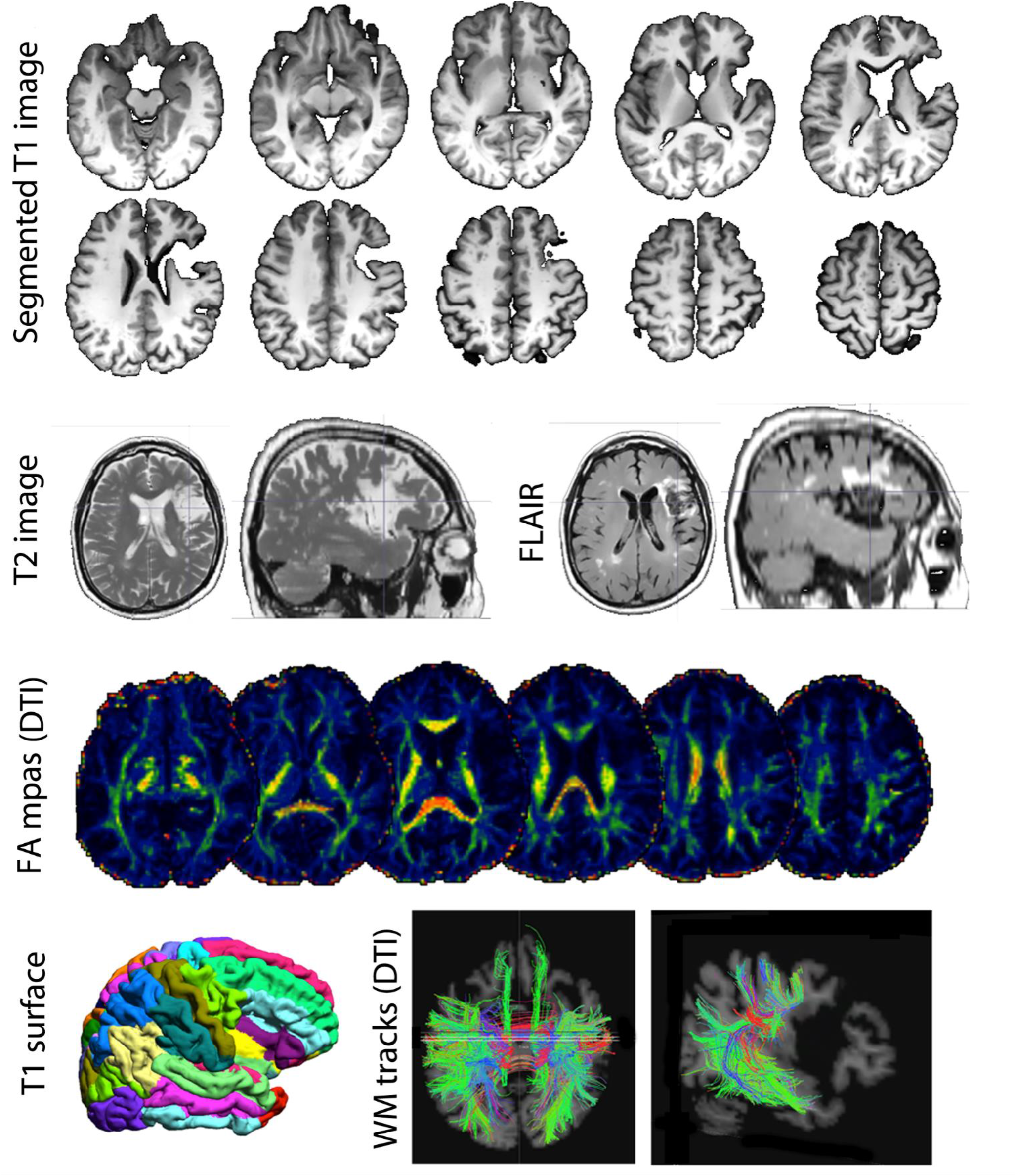
Structural imaging. At the top are shown the segmented T1 image, and corresponding T2 and FLAIR images, illustrating the location and size of the lesion. The bottom part of the figure show reconstructed images from the T1 (plial surface) and from the DTI data (fractional anisotropy (FA) maps and white matter (WM) tracks).

#### 3.2.2 Language assessment

Language was assessed longitudinally using The Western Aphasia Battery (WAB) (Kertesz, 1982) at the acute (1-6 weeks post stroke), sub-acute (2-6 months post stroke) and chronic (7-11 months post stroke) stages. At the acute and sub-acute time points CF was classified as ‘anomic’ (Aphasia Quotient of 79.9 and 87.8 respectively). At the chronic time point, CF obtained an Aphasia Quotient of 96.4 and was no longer classified as aphasic. Residual deficits were mild word-finding difficulties (fluency score = 9/10) and written output difficulties (written score = 91/100). His written output was characterised by syntactical errors, occasional phonemic orthographic output errors and cognitive demand errors. (Full breakdown of results are presented in Table A.4.)

To explore the exact breakdown of CF's language selected subtests of the Psycholinguistic Assessment of Language Processing in Aphasia (PALPA, Kay, Lesser, & Coltheart, 1992), an experimental battery of semantic assessments and an informal auditory phonological processing assessment were carried out and mapped onto an adapted cognitive-neuropsychological model (Figure 4).

**Figuer 4:**
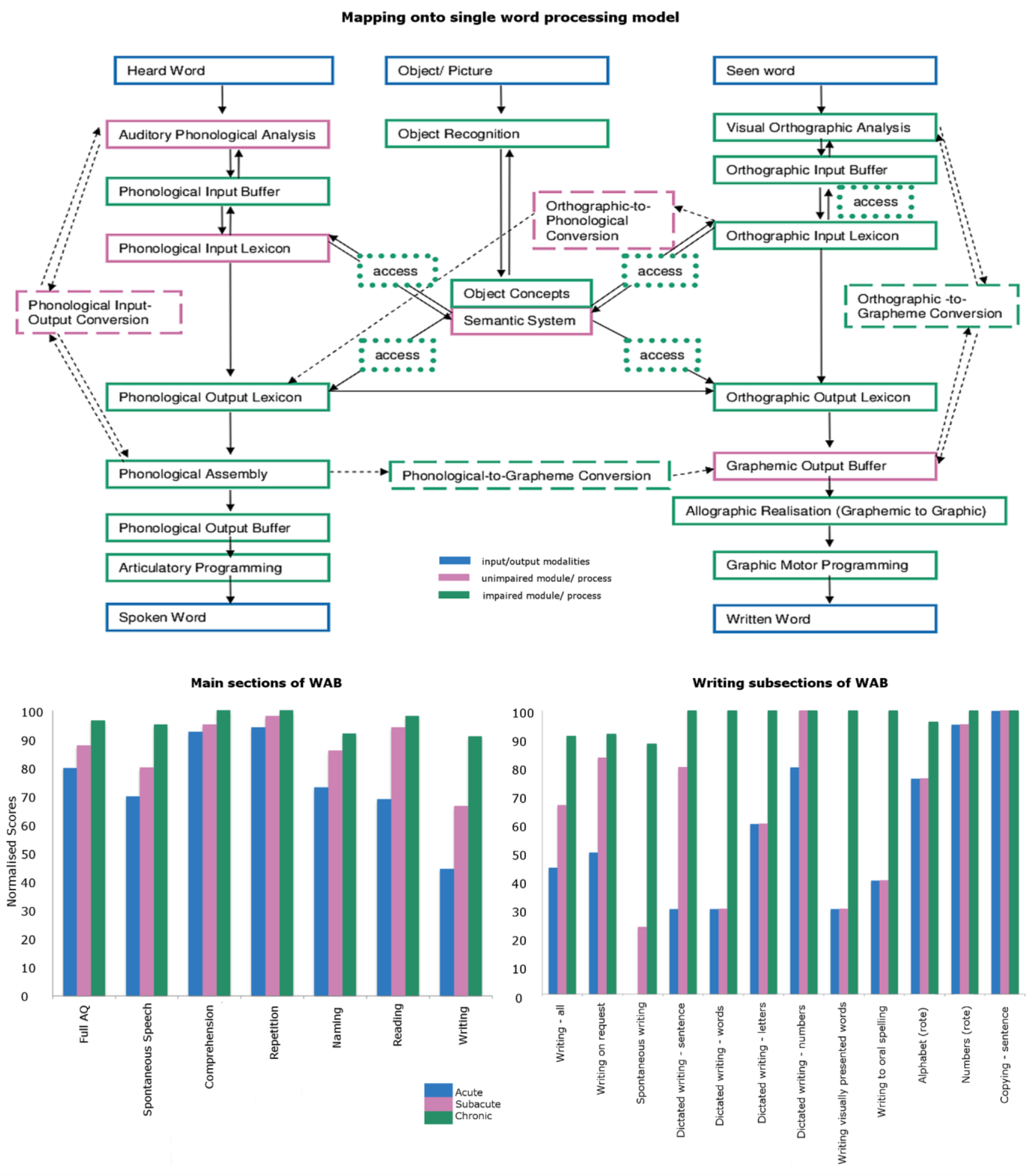
CF’s behavioural abilities at chronic time-point mapped onto a cognitive neuropsychological single word processing model (adapted from whitworth (2008) and Ellis (2004)). Blue: input and output modalities; Green unimpaired module (solid/dashed box)/ process (dotted box); Pink; impaired module (solid/dashed box)/process (dotted box). At the bottom, bar-graphs show CF’s normalised scores on the main sections and writing subsections of WAB at acute, sub-acute and chronic time.

<u>Phonological processing</u>, assessed using an informal auditory discrimination task at the acute/ subacute phase, showed an impaired distinction between /m/ and /n/; /k/ and /g/; /s/ and /z/. CF was, however, able to distinguish the following phoneme pairs: /p/ and /b/; /t/ and /d/; /⊖/ and /Φ/; /∫/ and /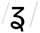/; /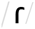/ and /l/. At the word level (assessed at the chronic phase), phoneme perception was also impaired with scores on PALPA 1 (non-word minimal pairs discrimination) showing deficits for both 'same’ and ‘different’ items, and scores on the PALPA 2 (real word minimal pairs discrimination) (Kay, Lesser, & Coltheart, 1992)(Kay, Lesser, & Coltheart, 1992)(Kay, Lesser, & Coltheart, 1992)(Kay, Lesser, & Coltheart, 1992)(Kay et al., 1992)(Kay et al., 1992)showing deficits for the 'same’ items but not for ‘different’ items. Together these results suggest that CF has impairment in auditory phonological analysis, which did not resolve between the acute and chronic phase. Assessment of <u>auditory lexical decision</u> using PALPA 5 at the chronic phase, showed no impairment with real word decisions but outside of the normal range of performance for non-words, indicating a breakdown in the phonological input lexicon. <u>Auditory input and spoken output</u> were assessed using PALPA 8 (non-word repetition) and PALPA 13 (digit span) at the chronic phase. CF showed an overall ability to repeat non-words with a few errors when increasing syllable length. His auditory digit span was unimpaired for both digit repetition and digit matching, with no length effects; which indicates that non-words repetition errors were caused by phonological output buffer impairment. <u>Auditory Comprehension</u> was assessed using subsections of the WAB at both the acute and chronic phases. Comprehension of single words and short phrases were within normal range at both stages. Comprehension of sentences was impaired at the acute phase, but was no longer impaired by the chronic stage.

Functional MRI revealed a right lateralized pattern of activation (Figure 3) for the verb generation and the word repetition tasks (based on lateralization curves), that was even stronger than the one observed for the 3 left handed control subjects tested (controls tested: right-handed n=7; left-handed n=3, (Gorgolewski, Storkey, Bastin, & Pernet, 2012). The verb generation tasks elicited activations in the right mid frontal gyrus and right posterior temporal cortex, almost mirroring the left frontal and temporal activations observed in controls. The word repetition task showed bilateral activations over the primary auditory cortices, but without extending posteriorly to Wernicke's area; instead strong right secondary auditory cortex activations were observed in CF.

**Figure 3:**
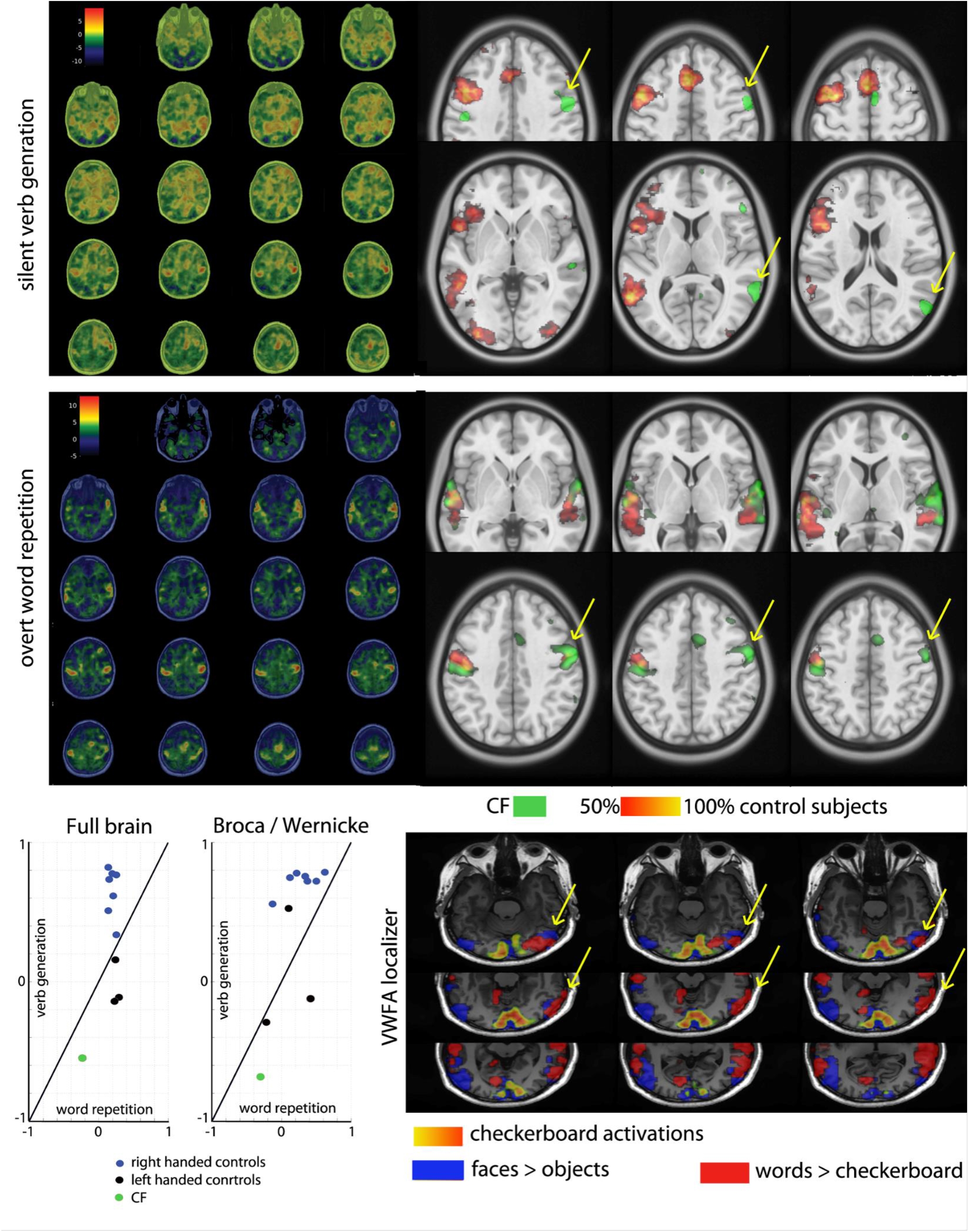
Functional Imaging results. On the left side (top and middle rows) are the unthresholded fMRI maps of CF for the verb generation task (visual input) and the overt word repetition task (auditory input). At the bottom (left) are plots showing the lateralization indices obtained in each task for CF compared to a control group of 10 subjects. The overlap of the single subject maps, projected into the standard space, can be seen on the right side (top and middle rows). The bottom right hand side shows the thresholded contrast maps for checkerboard (V1), faces (FFA) and words (VWFA).

<u>Written input (reading) and spoken output</u> (i.e. single word reading) was assessed by the WAB (word reading), PALPA 8 and PALPA 36 (both non-word reading), at the chronic stage. Performances on the WAB were within normal range, indicating intact lexical reading route. In contrast, non-word reading was impaired, with a length effect whereby non-words containing more than 2 syllables/ 4 letters were not read. This suggests an inability to hold long non-words in or the phonological output buffer and/or the orthographic-phonological conversion module.

The Visual Word Form Area was mapped using a passive localizer contrasting high frequency words to checkerboards (Cohen et al., 2000b; Dehaene & Cohen, 2011; Price & Devlin, 2011). Results show right lateralized activations (LI -0.35) over the right homologue of the Visual Word Form Area (Cohen et al., 2000a). During the same localizer, the FFA was mapped contrasting faces with objects, showing co-localization in the right hemisphere (LI = -0.2; Figure 3).

<u>Object input and written/spoken output</u> were tested at the chronic stage using PALPA 54 (picture to written and spoken outputs). CF did not have any difficulties with single word expression from pictorial inputs although he did make a few semantically related errors. <u>Object input and semantics:</u> CF was unimpaired in all of the semantic processing (noun, verb, syntagmatic and sounds) assessments, showing no central semantic system impairment.

#### 3.2.3 Dysgraphia

At the acute time point CF showed severely impaired written output on WAB (written score = 44.5/100). His written output improved at the sub-acute time point (written score = 66.5/100), although still showed the same pattern of errors as in the acute phase. At the chronic time point CF's written output was within normal limits, but was characterised by some syntactical errors, occasional phonemic orthographic output errors and cognitive demand errors (Figure 4; **Error! Reference source not found.**).

Written output was further examined at both the acute and chronic stages using PALPA 39 (words of varying letter length), PALPA 40 (words with varying imageability and frequency) and PALPA 45 (non-words dictation). At the acute stage, a severe dysgraphia was observed with an effect of word length, imageability and frequency and an inability to spell non-words. At the chronic stage, performances were within or close to normal for word spelling with errors but still showed a mild letter length effect and imageability effect, and no frequency effect. Despite performance recovery for words, non-word spelling was still severely impaired, indicative of a dysgraphia with a breakdown in phoneme/grapheme correspondence and at the level of the graphemic output buffer. An important observation is the marked difference between CF's spoken spelling and his written spelling at both time points - CF tended to spell the words dictated to him aloud in the correct form, at the same time as writing them down in an incorrect form. This dissociation clearly shows that CF's written output system is selectively impaired, with the spoken output system remaining intact, supporting impairment in phoneme/grapheme correspondence.

Interestingly, all three major areas involved in the ‘writing system’ (SFG, SMG and intra-parietal sulcus, 39) were physically intact. The lesion affects frontal areas inferior (lateral IFG) to the ‘writing system’ (SFG 39). The IFG and the SMG are both involved in grapheme/phoneme correspondence (Mei et al., 2014) and lesions of the fibers linking these regions could explain some of the observed deficits.

#### 3.2.4 Assessment of other cognitive abilities

CF presented with variable memory abilities suggesting external influencing factors such as fatigue, mood and stress/anxiety. CF performances’ were in the normal range for visuo-spatial skills and executive functioning. He did not show any signs of central apraxia even though there was damage to the MFG, showing that CF's praxis system was not damaged by the lesion (apraxias were split into ‘central’ and ‘peripheral’ as detailed in the methodology section). CF did not show any signs of visual neglect either, possibly because the lesion did not damage his parietal lobe, remaining more anterior. Acalculia screening showed an impairment when carrying out simple additions and deletions (Table A.1). These difficulties corresponded to numerical length, suggesting impairment in CF's output buffer rather than numeracy itself. He performed poorly on a specialist knowledge retention task (Figure A.1), prompting further, detailed, analysis of the results of this informal assessment to determine whether this was a similar presentation to those with semantic dementia and associated aphasia (Hirono et al., 2000; Graham, Lambon Ralph, & Hodges, 1999; Graham, Patterson, Pratt, & Hodges, 1999; Omar, Hailstone, Warren, Crutch, & Warren, 2010; Robinson, Rossor, & Cipolotti, 1999) or due to other associated impairments. On examination it was apparent that his difficulties were in word-finding and formulation of sentences, and not due to the actual retention of expert semantic knowledge (*Figure 4* & *Figure 5*). (Table A.5 shows cognitive profiles in full).

**Figure 5:**
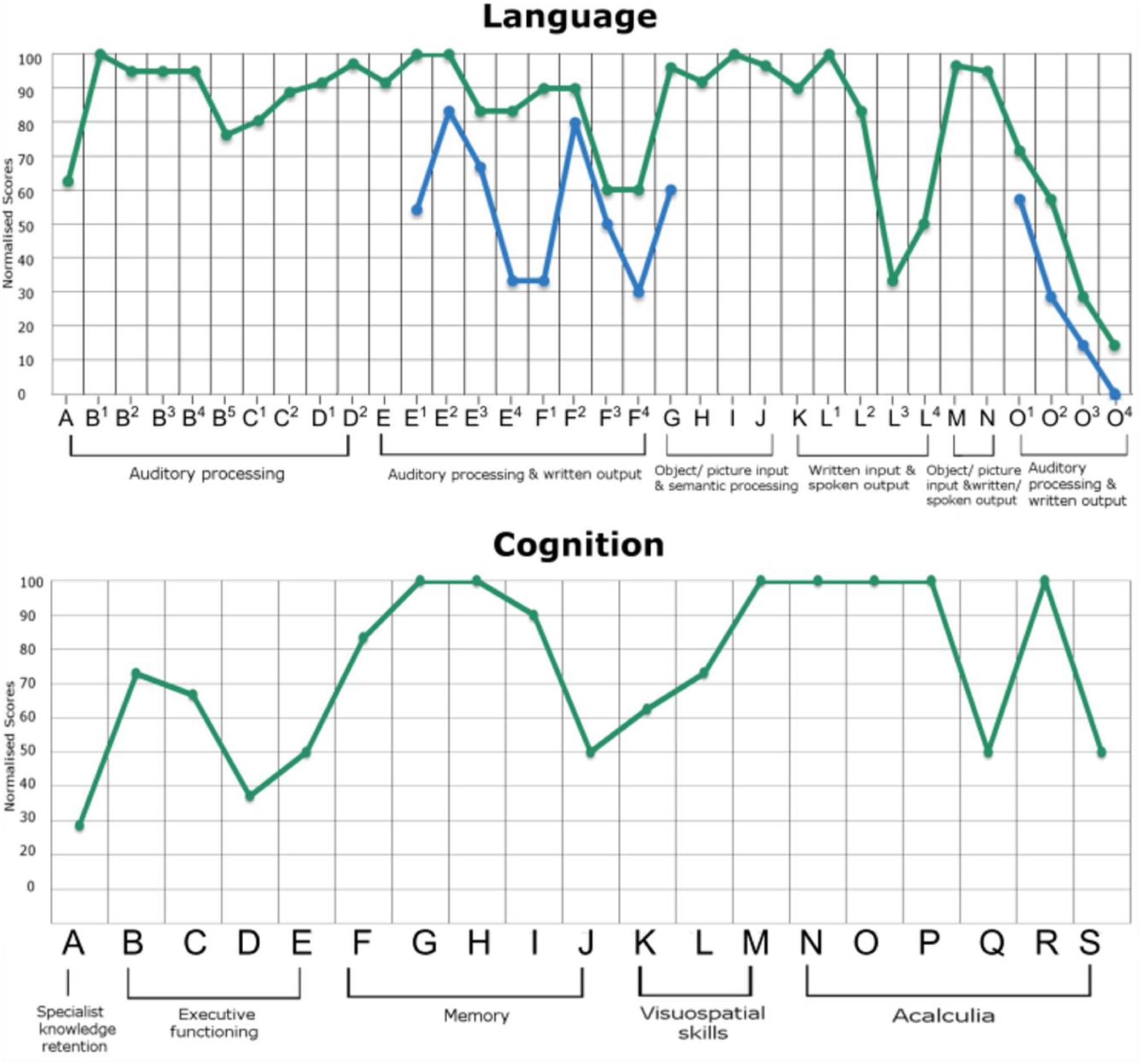
CF's language and cognitive profiles (normalised scores). For language (top), normalized scores are presented for phonological discrimination (A), PALPA 5 (high/low imageability, high/low frequency: B1, B2, B3, B4; non-words: B5), PALPA 1 (same/different judgements: C1, C2), PALPA 2 (same/different judgements: D1, D2), PALPA 39 (total: E; 3,4,5,6 letters: E1, E2, E3, E4), PALPA 40 (high/low imageability, high/low frequency: F1, F2, F3, F4), PPT (G), KDT (H), TTT (I), SPMT (J), PALPA 8 (K), PALPA 36 (3, 4, 5, 6 letters: L1, L2, L3, L4), PALPA 54 (written output: M; spoken output: N), PALPA 45 (3, 4, 5, 6 letters: O1, O2, O3, O4). For the other cognitive domains (bottom), normalized scores are presented for expertise retention (A), BADS (modified 6 elements: B; temporal judgement: C; key search: D); WAIS IV matrix reasoning (E); WMS II Faces I and II (visual immediate memory: F; visual delayed memory: G); BMIBP (total: H; speed of information processing: I), Rey-Osterrieth Complex figure (delayed recall: J; copying: K); VOSP (incomplete letters: L; dot counting: M); numeracy (counting forwards: N; counting backwards: O; size decisions: P; simple addition: Q; simple multiplications: R; simple deletions: S).

## 4. Discussion and conclusions

Using Mariën et al’s (2004) algorithm, we can conclude that CF is a reliable case of vascular crossed aphasia: he is a right-handed patient with no left-handedness in his family, showing morphological integrity of the left-hemisphere and no brain damage or seizures in childhood. CF shows clear-cut evidence of language disorder (anomic aphasia) with word-finding difficulties related to auditory phonological analysis. Functional MRI analyses show right hemispheric lateralization of language functions (reading, repetition, generation), which concurs with results from the literature review. Central apraxia, visual field neglect and acalculia were significantly associated with CA, which can only occur if they are right lateralized (as in most individuals) along with language, including reading and writing systems, which explain the association with agraphia.

CF presented also strong dysgraphia which we believe is a pre-existing, developmental disorder, not formally diagnosed due to lack of knowledge and diagnosis available at this time (Swanson, Harris, & Graham, 2013). Developmental dysgraphia can be defined as “a specific learning disorder […] an impairment in written expression” (American Psychiatric Association, 2013) that causes problems in handwriting only, spelling only, or both handwriting and spelling (Berninger, Abbott, Thomson, & Raskind, 2001). CF reported problems from a ‘young age’ *only* with spelling - i.e. he did not have any other developmental or medical conditions, and he presents with a breakdown in his graphemic output buffer (see figure 3 - final row), leading to difficulty spelling non-words and effects of word length, imageability and frequency - impairments involving ‘writing specific’ processes (Planton, Jucla, Roux, & Demonet, 2013). CF’s dysgraphia can further be classified as ‘dyslexic aphasic dysgraphia’ i.e. a language disorder mainly characterised by a writing impairments consisting of “mis-spellings with reversals, omissions, inversions and substitutions non words and paragraphic errors” (Gubbay & De Klerk, 1995) consistent with an impairment in phoneme/grapheme correspondence. Impaired non-lexical reading was also present, also suggesting the presence of mild phonological dyslexia, which concurs with the idea of a pre-morbid impairment in phoneme/grapheme correspondence. Advances in imaging techniques have allowed the investigation of the neurological basis of dyslexia (Shaywitz et al., 1998; Shaywitz et al., 2002; Shaywitz et al., 2003; Shaywitz, Mody, & Shaywitz, 2006; Cao, Bitan, Chou, Burman, & Booth, 2006; Habib, 2000). It has been shown that deficits in the left inferior frontal gyrus, left inferior parietal lobule (i.e. supramarginal gyrus & angular gyrus: Singh-Curry & Husain, 2009), and mid-ventral temporal cortex (i.e. fusiform gyrus, parahippocampal gyrus, lingual gyri & inferior temporal gyri: Haxby et al., 2001) are associated with developmental dyslexia in dextrals. Shaywitz (2002; 2003) has further suggested that such left hemisphere disruptions are compensated for by recruitment of the right hemisphere, supporting the theory that developmental disorders can be an underlying cause of crossed aphasia.

Putting this deficit back into the general context of CA, we can postulate that CF had left hemisphere defects causing the dysgraphia and dyslexia and causing a right hemispheric language shift. As recent evidence suggests that lateralisation shift occurs not only with large lesions, but also small focal lesions or dysfunction of neural networks (Guerreiro, Castrocaldas, & Martins, 1995; Lazar et al., 2000; Maesto et al., 2004; Staudt et al., 2001; Kurthen, Linke, Elger, & Schramm, 1992), we conclude that CA can be caused by a congenital dysfunction within the left reading/writing systems, and not just the left auditory/spoken system (Bakar, Kirshner, & Wertz, 1996; Bhatnagar, Imes, Buckingham, & Puglishi-Creegan, 2006; Cappa et al., 1993). In addition, since (i) only 1 out of the 57 cases identified in the review could be conclusively attributed to a genetic basis (Cohen, Grony, Hermine, Gray, & Degos, 1993), (ii) bi-hemispheric representation was suggested in 10 of the remaining 56 cases (Bakar, Kirshner, & Wertz, 1996; Cappa et al., 1993; Giovagnoli, 1993; Habib, Joanette, Ali-Cherif, & Poncet, 1983; Ishizaki et al., 2012; Paghera, Marien, & Vignolo, 2003; Paparounas, Eftaxias, & Akritidis, 2002), and (iii) when tested, dysgraphia co-occurred in >60% of CA cases, it is conceivable that developmental disorders cause a total or partial right lateralisation shift in language functioning, at least in some cases.

## Acknowledgements

AJ is funded by SINAPSE under the SPIRIT scheme (Scottish Funding Council).

## Conflicts of Interest

The authors declare that there are no conflicts of interest.

